# Utility of AlphaMissense predictions in Asparagine Synthetase deficiency variant classification

**DOI:** 10.1101/2023.10.30.564808

**Authors:** Stephen J. Staklinski, Armin Scheben, Adam Siepel, Michael S. Kilberg

## Abstract

AlphaMissense is a recently developed method that is designed to classify missense variants into pathogenic, benign, or ambiguous categories across the entire human proteome. Asparagine Synthetase Deficiency (ASNSD) is a developmental disorder associated with severe symptoms, including congenital microcephaly, seizures, and premature death. Diagnosing ASNSD relies on identifying mutations in the asparagine synthetase (ASNS) gene through DNA sequencing and determining whether these variants are pathogenic or benign. Pathogenic ASNS variants are predicted to disrupt the protein’s structure and/or function, leading to asparagine depletion within cells and inhibition of cell growth. AlphaMissense offers a promising solution for the rapid classification of ASNS variants established by DNA sequencing and provides a community resource of pathogenicity scores and classifications for newly diagnosed ASNSD patients. Here, we assessed AlphaMissense’s utility in ASNSD by benchmarking it against known critical residues in ASNS and evaluating its performance against a list of previously reported ASNSD-associated variants. We also present a pipeline to calculate AlphaMissense scores for any protein in the UniProt database. AlphaMissense accurately attributed a high average pathogenicity score to known critical residues within the two ASNS active sites and the connecting intramolecular tunnel. The program successfully categorized 78.9% of known ASNSD-associated missense variants as pathogenic. The remaining variants were primarily labeled as ambiguous, with a smaller proportion classified as benign. This study underscores the potential role of AlphaMissense in classifying ASNS variants in suspected cases of ASNSD, potentially providing clarity to patients and their families grappling with ongoing diagnostic uncertainty.

## Introduction

Asparagine Synthetase deficiency (ASNSD) is a rare developmental disorder, caused by biallelic mutations in the Asparagine Synthetase (ASNS) gene, initially documented in 2013^1^. Children with ASNSD often exhibit clinical symptoms, including intractable seizures, congenital microcephaly, physical and mental developmental delay, continued brain atrophy, and premature mortality^2–4^. Currently, diagnosing ASNSD remains challenging, primarily relying on DNA sequencing^2^, and classifying the variants caused by identified mutations as benign or pathogenic is not straightforward. Cases of ASNSD have been primarily reported through clinical case studies, although a few studies initiated the investigation of the suspected ASNS variants using laboratory-based techniques^1,5,6^. When it is not possible to conduct time-consuming and expensive laboratory experiments on suspected variants, the classification may depend on a general understanding of ASNS’s structure and function, along with computational variant effect prediction tools.

ASNS functions in an ATP-dependent manner to synthesize asparagine and glutamate from the substrates aspartate and glutamine. The ASNS protein contains 561 amino acids organized into two domains. The N-terminal domain active site exhibits glutaminase activity, generating an ammonia group that traverses an intramolecular tunnel to the C-terminal domain active site^7^. Within the C-terminal domain, activation of aspartate by ATP results in a β-aspartyl-AMP intermediate. The intermediate combines with the ammonia to produce asparagine, with the byproducts of glutamate and AMP^8^. Mutations within the ASNS gene commonly present as missense mutations, which involve the substitution of a single amino acid, or frameshift mutations, often resulting in premature truncation of the protein (Table 1). These mutations can give rise to pathogenic variations by diminishing the functionality of the ASNS protein, either through a reduction in enzymatic activity or the destabilization of its structure^9–12^. Depending on the cell’s ability to acquire asparagine from the extracellular milieu, pathogenic ASNS variants can lead to a disruption of global metabolism due to intracellular starvation for asparagine, inhibiting cell growth^13^. Palmer et al.^5^ provided the initial recognition of this phenomenon as the molecular underpinning of ASNSD-associated neurological deficiency. The brain is particularly reliant on *de novo* asparagine synthesis due to the active export of asparagine across the blood-brain barrier^14^. This explanation is consistent with the impact of ASNSD on individual tissues, of which the brain is predominantly affected^2^. Therefore, the diagnosis of ASNSD hinges on distinguishing newly identified mutations as either benign or pathogenic variants affecting the ability of ASNS to synthesize asparagine in the brain.

**Table 1:**
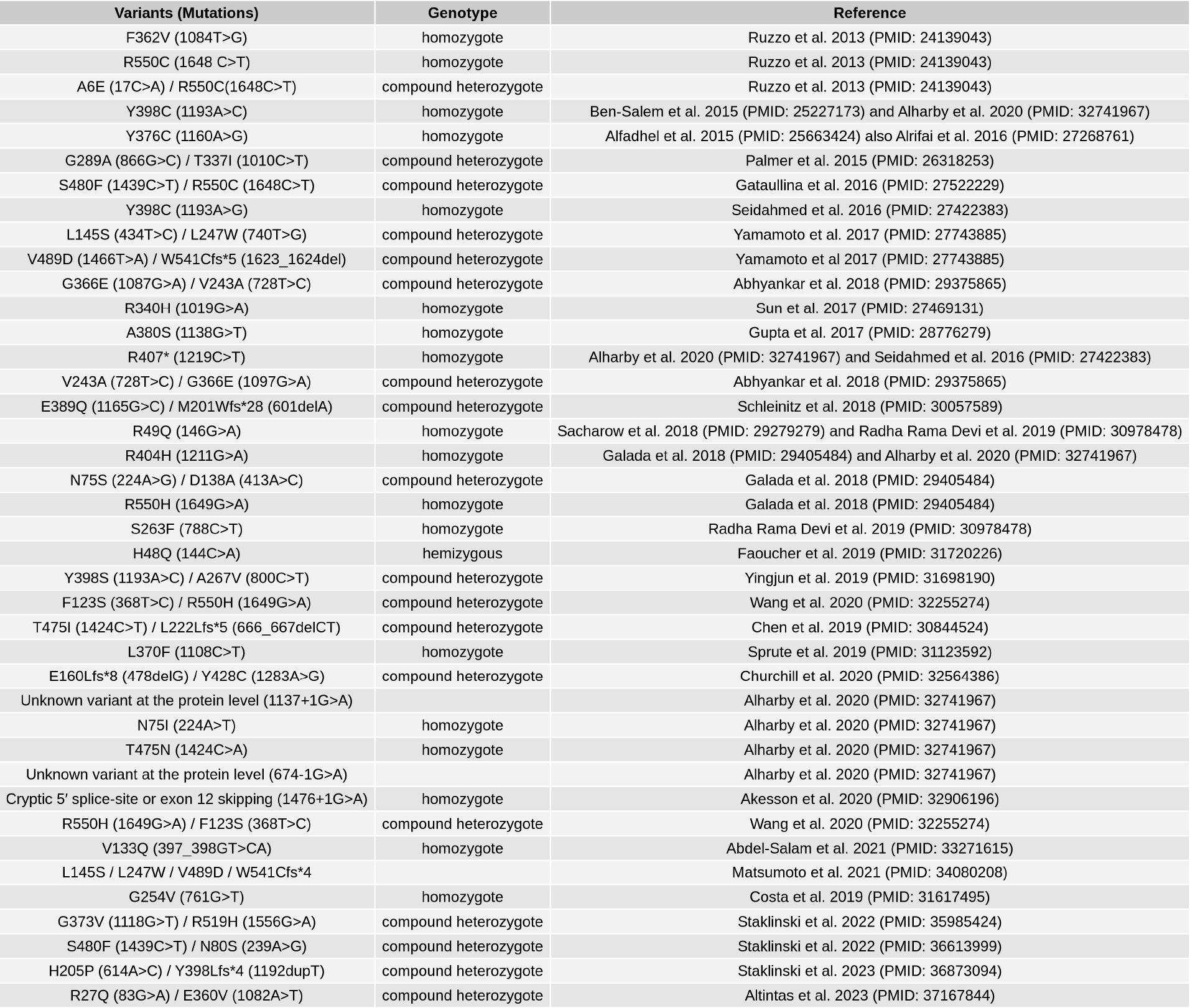
Summary of previously reported ASNSD variants from a literature review. The first column contains the ASNS protein variant(s) (UniProt P08243) with DNA mutations in parentheses following the variant notation. A single variant was included for homozygous patients, whereas two values were included for compound heterozygote patients, separated by a forward slash. The second column classifies the genotype into homozygous, compound heterozygous, or hemizygous categories. The third column provides the reference from which the data were obtained and includes the PubMed ID (PMID) for easy access.

| Variants (Mutations) | Genotype | Reference |
| --- | --- | --- |
| F362V (1084T>G) | homozygote | Ruzzo et al. 2013 (PMID: 24139043) |
| R550C (1648 C>T) | homozygote | Ruzzo et al. 2013 (PMID: 24139043) |
| A6E (17C>A) / R550C(1648C>T) | compound heterozygote | Ruzzo et al. 2013 (PMID: 24139043) |
| Y398C (1193A>C) | homozygote | Ben-Salem et al. 2015 (PMID: 25227173) and Alharby et al. 2020 (PMID: 32741967) |
| Y376C (1160A>G) | homozygote | Alfadhel et al. 2015 (PMID: 25663424) also Alrifai et al. 2016 (PMID: 27268761) |
| G289A (866G>C) / T337I (1010C>T) | compound heterozygote | Palmer et al. 2015 (PMID: 26318253) |
| S480F (1439C>T) / R550C (1648C>T) | compound heterozygote | Gataullina et al. 2016 (PMID: 27522229) |
| Y398C (1193A>G) | homozygote | Seidahmed et al. 2016 (PMID: 27422383) |
| L145S (434T>C) / L247W (740T>G) | compound heterozygote | Yamamoto et al. 2017 (PMID: 27743885) |
| V489D (1466T>A) / W541Cfs*5 (1623_1624del) | compound heterozygote | Yamamoto et al 2017 (PMID: 27743885) |
| G366E (1087G>A) / V243A (728T>C) | compound heterozygote | Abhyankar et al. 2018 (PMID: 29375865) |
| R340H (1019G>A) | homozygote | Sun et al. 2017 (PMID: 27469131) |
| A380S (1138G>T) | homozygote | Gupta et al. 2017 (PMID: 28776279) |
| R407* (1219C>T) | homozygote | Alharby et al. 2020 (PMID: 32741967) and Seidahmed et al. 2016 (PMID: 27422383) |
| V243A (728T>C) / G366E (1097G>A) | compound heterozygote | Abhyankar et al. 2018 (PMID: 29375865) |
| E389Q (1165G>C) / M201Wfs*28 (601delA) | compound heterozygote | Schleinitz et al. 2018 (PMID: 30057589) |
| R49Q (146G>A) | homozygote | Sacharow et al. 2018 (PMID: 29279279) and Radha Rama Devi et al. 2019 (PMID: 30978478) |
| R404H (1211G>A) | homozygote | Galada et al. 2018 (PMID: 29405484) and Alharby et al. 2020 (PMID: 32741967) |
| N75S (224A>G) / D138A (413A>C) | compound heterozygote | Galada et al. 2018 (PMID: 29405484) |
| R550H (1649G>A) | homozygote | Galada et al. 2018 (PMID: 29405484) |
| S263F (788C>T) | homozygote | Radha Rama Devi et al. 2019 (PMID: 30978478) |
| H48Q (144C>A) | hemizygous | Faoucher et al. 2019 (PMID: 31720226) |
| Y398S (1193A>C) / A267V (800C>T) | compound heterozygote | Yingjun et al. 2019 (PMID: 31698190) |
| F123S (368T>C) / R550H (1649G>A) | compound heterozygote | Wang et al. 2020 (PMID: 32255274) |
| T475I (1424C>T) / L222Lfs*5 (666_667delCT) | compound heterozygote | Chen et al. 2019 (PMID: 30844524) |
| L370F (1108C>T) | homozygote | Sprute et al. 2019 (PMID: 31123592) |
| E160Lfs*8 (478delG) / Y428C (1283A>G) | compound heterozygote | Churchill et al. 2020 (PMID: 32564386) |
| Unknown variant at the protein level (1137+1G>A) |  | Alharby et al. 2020 (PMID: 32741967) |
| N75I (224A>T) | homozygote | Alharby et al. 2020 (PMID: 32741967) |
| T475N (1424C>A) | homozygote | Alharby et al. 2020 (PMID: 32741967) |
| Unknown variant at the protein level (674-1G>A) |  | Alharby et al. 2020 (PMID: 32741967) |
| Cryptic 5' splice-site or exon 12 skipping (1476+1G>A) | homozygote | Akesson et al. 2020 (PMID: 32906196) |
| R550H (1649G>A) / F123S (368T>C) | compound heterozygote | Wang et al. 2020 (PMID: 32255274) |
| V133Q (397_398GT>CA) | homozygote | Abdel-Salam et al. 2021 (PMID: 33271615) |
| L145S / L247W / V489D / W541Cfs*4 |  | Matsumoto et al. 2021 (PMID: 34080208) |
| G254V (761G>T) | homozygote | Costa et al. 2019 (PMID: 31617495) |
| G373V (1118G>T) / R519H (1556G>A) | compound heterozygote | Staklinski et al. 2022 (PMID: 35985424) |
| S480F (1439C>T) / N80S (239A>G) | compound heterozygote | Staklinski et al. 2022 (PMID: 36613999) |
| H205P (614A>C) / Y398Lfs*4 (1192dupT) | compound heterozygote | Staklinski et al. 2023 (PMID: 36873094) |
| R27Q (83G>A) / E360V (1082A>T) | compound heterozygote | Altintas et al. 2023 (PMID: 37167844) |

An important problem in human genetics is the classification of variants of unknown significance, particularly those with the potential to be loss of function variants^15^. Single variants can often be classified through biochemical assays, as is currently done when possible in ASNSD^9,10^. Multiplex assays of variant effect are promising high throughput variant classification experiments^16^, but the costs are often prohibitive and the approach is dependent on an accurate readout metric for a protein’s function. A multiplex assay of variant effect has not been reported for ASNS. As a result, the initial classification of potentially pathogenic ASNS variants relies on computational tools that commonly predict pathogenicity based on evolutionary features such as sequence constraint^17^.

These tools rely on nucleotide-level sequence conservation^18–20^, the co-variance of amino acids within a sequence^21^, integration of sequence conservation and functional data^22–24^, or ensemble approaches that use machine learning models trained on variant scores from multiple tools^25^. Most recently, AlphaMissense boasts an improved performance through an all-in-one method utilizing population variant frequency data, amino acid distribution, and structural context premised on an AlphaFold protein structure prediction system^26,27^. With the demonstrated importance of AlphaFold in recent years and the purported effective performance of AlphaMissense, it is plausible to expect that researchers will increasingly turn to AlphaMissense for predicting the effect of missense variants.

In this report, the performance of AlphaMissense in predicting the pathogenicity of ASNS missense variants was assessed by comparing its efficacy against a curated list of known ASNSD-associated missense variants. To enable universal AlphaMissense data visualization for any human protein with a UniProt ID, a user-friendly Snakemake pipeline was created. To validate AlphaMissense’s predictive capability with ASNS, critical residues within ASNS active sites and the intramolecular tunnel with established importance were employed as benchmarks. AlphaMissense effectively identified these critical residues as having a high pathogenicity score if replaced. It was noteworthy that the intramolecular tunnel residues recently discovered^28^ were supported by the AlphaMissense predictions of their importance within the ASNS structure. With known ASNSD missense variants, AlphaMissense categorized 78.9% of variants as pathogenic, with the majority of the remaining classifications falling into the ambiguous category. This benchmark study highlights the basic utility of AlphaMissense in classifying ASNS variants, albeit with inherent uncertainty in roughly 20% of classifications. To the best of our knowledge, this is the first study assessing the performance of a computational tool for categorizing variant effects in ASNSD. Additionally, this report marks one of the early efforts to utilize the amino acid substitution dataset from AlphaMissense to assess performance against known pathogenic variants within a particular gene context, beyond those examples that were initially presented by the authors of AlphaMissense.

## Results

### Snakemake pipeline for the analysis of AlphaMissense pathogenicity scores for any human UniProt ID

There is often a gap between computational biologists who develop advanced tools for predicting the impact of protein variants and clinicians or experimental biologists who need to interpret new protein variants, but may not be proficient in computational techniques. To address this, we developed a Snakemake^29^ pipeline that allows researchers to easily access and analyze the AlphaMissense pathogenicity prediction resources for the entire human proteome. The pipeline comes with installation documentation to assist researchers with varying levels of computational skills. The Snakemake pipeline automatically downloads the raw AlphaMissense dataset, and the user can specify a UniProt ID corresponding to the protein they are interested in probing. The pipeline then extracts data specific to the provided UniProt ID from the complete AlphaMissense dataset. The average pathogenicity score for all 19 possible variants at each amino acid position is computed and used to generate a user-friendly plot displaying these average scores for the target protein. The AlphaFold prediction^27^ for the input UniProt ID is automatically retrieved and the PDB file is rewritten so that the beta factors represent the average AlphaMissense pathogenicity score per position. The user can then import this modified PDB into protein visualization software to color the protein structure based on the average pathogenicity score at each residue. Lastly, a heatmap of the scores for all possible variants in all positions is plotted. The heatmap and modified PDB file are directly modeled after the data visualization used in the original AlphaMissense paper^26^. The plot of average pathogenicity scores for each residue is an additional visualization that we believe compliments the colored protein structure well.

### Average pathogenicity score by amino acid aligns with known ASNS critical residues and provides further evidence for an intramolecular tunnel

Research on the asparagine synthetase protein has spanned many years, investigating its structure, function, and regulation. Within ASNS, specific residues play crucial roles at either the N or C terminal catalytic sites^8^. Evaluating the AlphaMissense pathogenicity score for these well-established critical residues can help gauge the utility of AlphaMissense in relation to ASNS enzymatic activity. Additionally, it has long been suggested that ASNS possesses an intramolecular tunnel responsible for transporting ammonia, a reaction intermediate^7^. Recently, these tunnel-related residues were identified through 3D variability analysis of an ASNS structure and complementary molecular docking simulations^28^. Upon superimposing the human ASNS crystal structure (PDB 6GQ3) with a per-residue color-coded representation based on the average AlphaMissense pathogenicity score, a visual pattern emerges. The highest pathogenicity scores are clustered in close proximity to the N and C terminal active sites, as well as at the interface connecting these two terminal regions. As anticipated, the pathogenicity scores are lower for the exterior residues of the protein (Fig. 1A). When plotting the average pathogenicity score per residue from AlphaMissense for ASNS, emphasizing the known N-terminal active site residues, C-terminal active site residues, and intramolecular tunnel residues, it is evident that AlphaMissense not only predicts the significance of known critical residues, but also strengthens the case for the newly identified intramolecular tunnel residues by predicting high pathogenicity scores upon their variation (Fig. 1B). These findings underscore the pivotal role these residues play in the protein’s function and help validate the utility of AlphaMissense using ASNS as a model.

**Figure 1:**
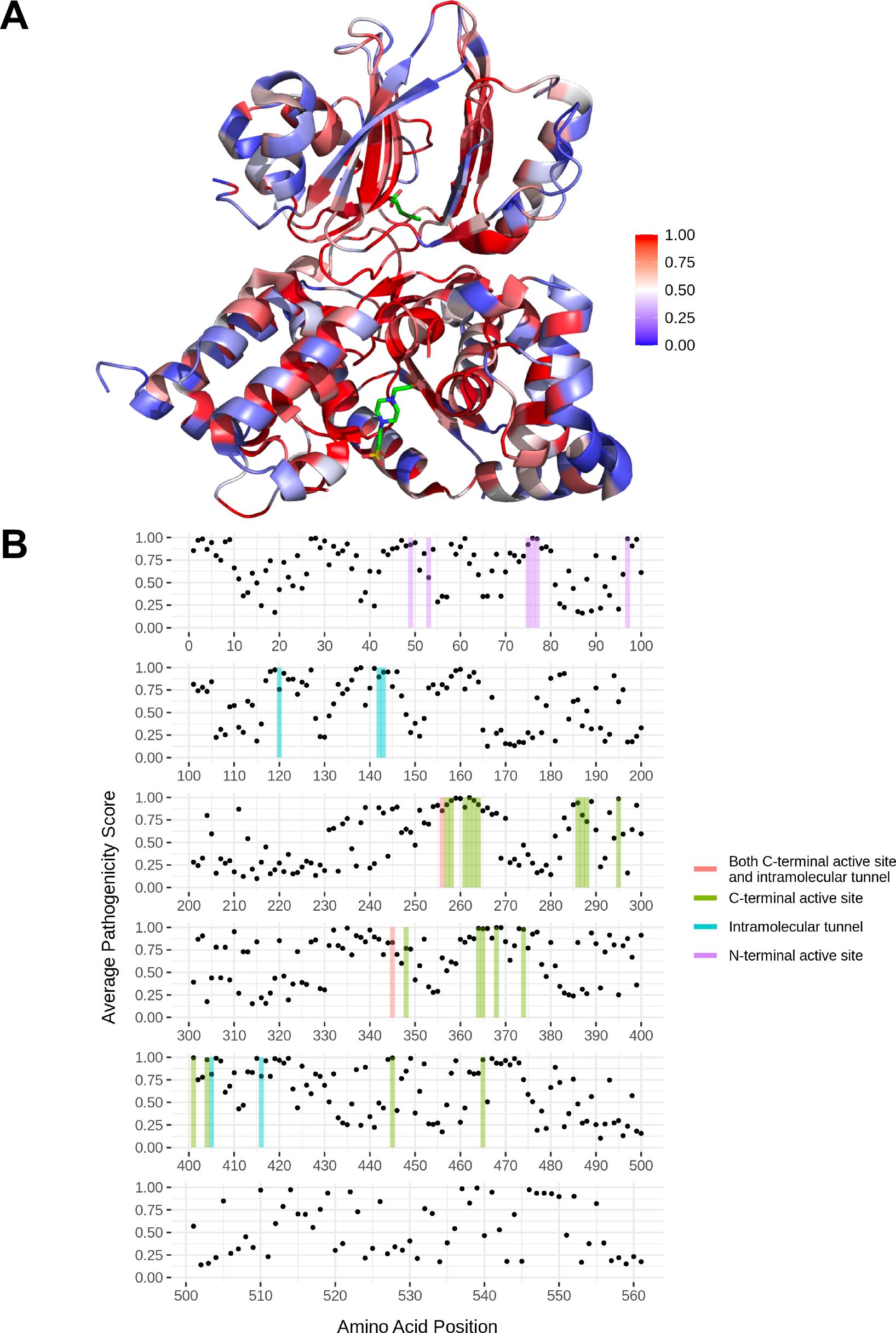
Average AlphaMissense pathogenicity score per amino acid in ASNS. (A) ASNS protein structure colored by average AlphaMissense pathogenicity score per amino acid. The N-terminal glutamine binding domain is shown at the top and the C-terminal ATP/aspartate binding domain is shown at the bottom. The molecules shown in green within each region were retained from the human ASNS (PDB 6GQ3) structure for a relative visual reference of binding pockets. The N-terminal site is bound to HEPES and the C-terminal site is bound to DON (6-diazo-5-oxo-L-norleucine). (B) Average pathogenicity score per amino acid plotted with an overlay of color shades to identify key residues in both the intramolecular tunnel and C-terminal active site (pink), the C-terminal active site only (green), the intramolecular tunnel only (blue), and the N-terminal active site only (purple).

### AlphaMissense accurately predicts the pathogenicity of known ASNSD variants

Diagnosing ASNSD relies on identifying mutations in both alleles of the ASNS gene through DNA sequencing^2^. However, determining if a mutation results in a deleterious protein variant can be challenging, often necessitating the use of predictive computational models. It is essential to assess the performance of AlphaMissense, a promising protein variant prediction model, on ASNSD variants that have been previously characterized. To achieve this comparison, it was necessary to compile a list of reported ASNSD variants, patient genotypes, and associated references (Table 1)^1,3,5,6,9–12,30–51^. The list could then be narrowed down to missense mutations causing substitution variants for which the AlphaMissense predictions could be obtained from the proteome-wide community resource on amino acid substitutions (Table 2). Amino acid positions harboring reported ASNSD-associated substitution variants were distinguished amongst AlphaMissense average pathogenicity scores for all amino acid positions. This analysis showed that ASNSD variants are distributed relatively uniformly across the entire ASNS protein sequence. However, there are localized patterns of ASNSD variants near regions with peaks of high average AlphaMissense pathogenicity scores (Fig. 2A). It is important to note that the average pathogenicity score for a position may not be representative of the score for a single variant. This discrepancy could be due to heterogeneity in the scores for one position, likely resulting from the biochemical properties of the substituted amino acid. In ASNS, this variation is evident for some positions through the display of all possible missense variant scores in a heatmap, from which ASNSD-associated variants can be located (Figure 2B). Analyzing 38 unique ASNSD-associated missense variants revealed that AlphaMissense performed reasonably well, correctly classifying 30 variants as pathogenic (78.9%), five as ambiguous (13.2%), and three as benign (7.9%) (Table 2). Notably, most misclassifications involved known ASNSD variants being categorized as ambiguous, indicating uncertainty to the user. Among the 8 known ASNSD variants predicted as ambiguous or benign, five (62.5%) were reported in compound heterozygous patients. Strikingly, although the objective of this study is not to predict new ASNSD-associated variants, there are 5,908 additional missense variants that are predicted to be pathogenic and distributed throughout the ASNS protein, covering 55.9% of all possible missense variants. While this performance does not allow one to definitively confirm the deleterious nature of novel ASNS missense variants in all cases, it does provide a valuable initial classification that is quicker and much more accessible than extensive enzymatic and cellular-based assays.

**Table 2:** AlphaMissense pathogenicity scores and predictions for reported ASNSD missense variants. Some missense variants were reported in multiple patients in the literature, but are represented here as one entry in the total list of unique variants.

| variant | score | prediction |
| --- | --- | --- |
| A267V | 0.5287 | ambiguous |
| N75S | 0.3977 | ambiguous |
| T475N | 0.4381 | ambiguous |
| Y398S | 0.3603 | ambiguous |
| Y428C | 0.4533 | ambiguous |
| A380S | 0.2278 | benign |
| N80S | 0.3009 | benign |
| Y398C | 0.1724 | benign |
| A6E | 0.9920 | pathogenic |
| D138A | 0.9972 | pathogenic |
| E389Q | 0.9253 | pathogenic |
| F123S | 0.9313 | pathogenic |
| F362V | 0.7241 | pathogenic |
| G254V | 0.9080 | pathogenic |
| G289A | 0.7090 | pathogenic |
| G366E | 0.9964 | pathogenic |
| G373V | 0.9973 | pathogenic |
| H205P | 0.6953 | pathogenic |
| H48Q | 0.8532 | pathogenic |
| L145S | 0.8879 | pathogenic |
| L247W | 0.6385 | pathogenic |
| L370F | 0.8516 | pathogenic |
| N75I | 0.9806 | pathogenic |
| R27Q | 0.9595 | pathogenic |
| R340H | 0.9178 | pathogenic |
| R404H | 0.9469 | pathogenic |
| R49Q | 0.7735 | pathogenic |
| R519H | 0.7756 | pathogenic |
| R550C | 0.7701 | pathogenic |
| R550H | 0.7667 | pathogenic |
| S263F | 0.9957 | pathogenic |
| S480F | 0.8733 | pathogenic |
| T337I | 0.8501 | pathogenic |
| T475I | 0.7093 | pathogenic |
| V133Q | 0.9830 | pathogenic |
| V243A | 0.7264 | pathogenic |
| V489D | 0.8666 | pathogenic |
| Y376C | 0.8623 | pathogenic |

**Figure 2:**
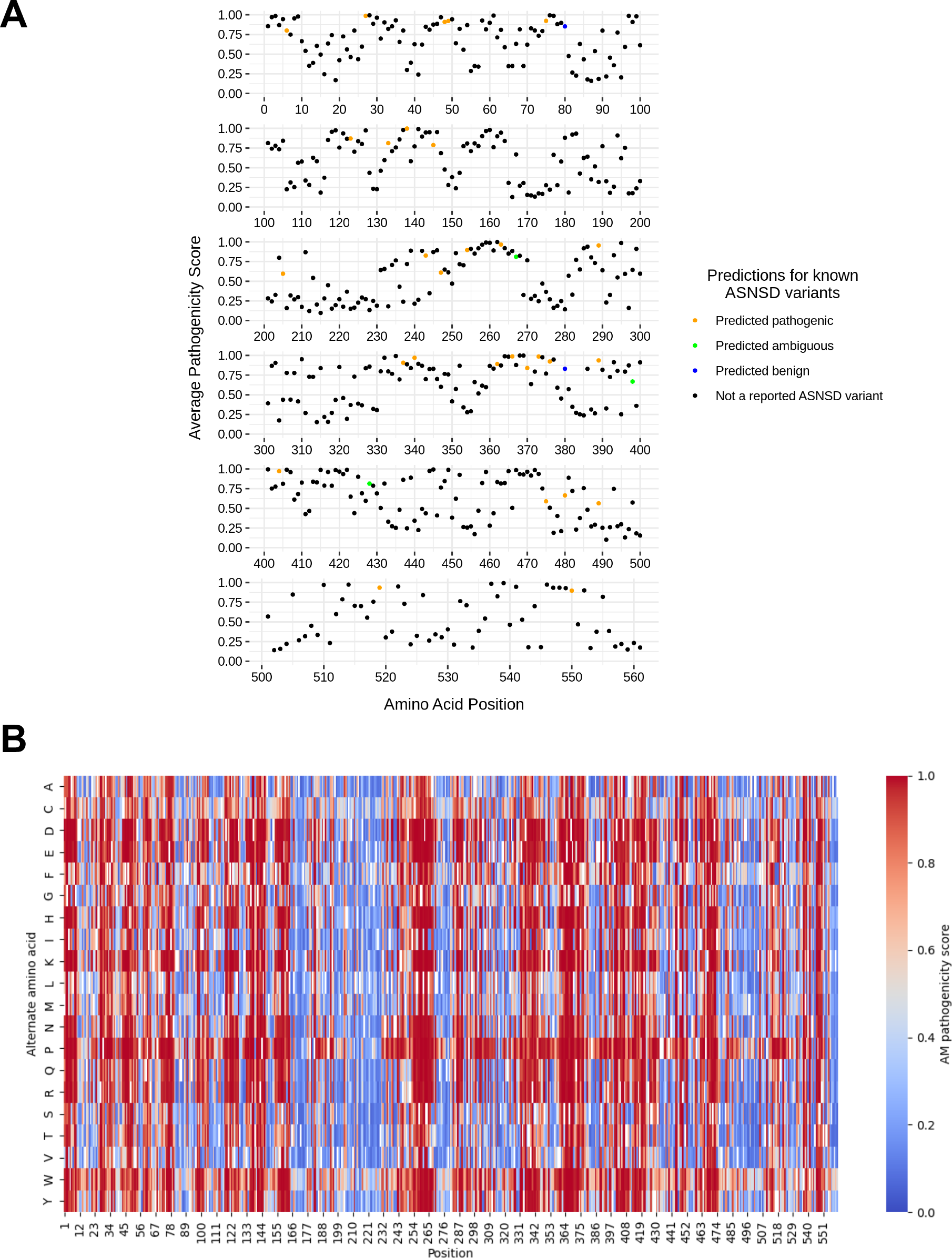
AlphaMissense pathogenicity scores for known ASNSD missense variants. (A) ASNS average AlphaMissense pathogenicity score by amino acid overlayed with the predictions for known ASNSD variants. The colors represent the classification of one ASNSD-associated variant and are not directly responsible for the score at which the point is placed as the score is an average of all possible 19 variants at that position. The amino acid positions with a known variant predicted by AlphaMissense to be pathogenic are shown in orange, ambiguous in green, and benign in blue. The amino acid positions without a published ASNSD variant are shown in black. Amino acid position 398 had two distinct ASNSD-associated variants with one classified as ambiguous and the other as benign, resulting in an ambiguous color for the plot. Amino acid positions 75 and 475 each had two ASNSD-associated variants, one with a pathogenic classification and the other with an ambiguous classification, resulting in a pathogenic coloring in the plot. (B) Heatmap of AlphaMissense pathogenicity scores for all possible missense variants at all positions within the 561 amino acid sequence for ASNS.

## Discussion

The classification of newly identified protein variants as either harmful or benign has long been a challenge in the field of biology. Various computational prediction methods have been developed, predominantly relying on evolutionary classification criteria, which assess sequence conservation (or its absence) across closely related species. Recently, a novel approach called AlphaMissense has expanded on this sequence conservation metric by incorporating metrics from other machine learning models as well as structural data obtained from pre-training with aspects of the groundbreaking work of AlphaFold, which accurately predicted the overall structure of all human proteins just a few years ago^26,27^. While AlphaMissense holds promise in terms of its performance when analyzed by proteome and genome-wide statistics, it is crucial to evaluate its applicability on specific proteins, particularly those for which variant classification plays an immediate role in a patient’s diagnosis.

A prime example is Asparagine Synthetase Deficiency (ASNSD), a disease for which a child’s diagnosis is based entirely on DNA sequencing and therefore, intimately linked to the identification of deleterious protein variants through the detection of mutations^2–4^. Precise variant classification is of utmost importance to ensure the correct treatment for the patient, whether it involves addressing ASNSD or seeking an alternative diagnosis more suitable in cases of a benign ASNS mutation. The present study was designed to assess the effectiveness of AlphaMissense in accurately predicting the pathogenicity of ASNS variants by benchmarking it against variants previously reported in cases of ASNSD.

To facilitate similar analyses for other researchers and clinicians studying Mendelian diseases or investigating individual proteins, we developed a straightforward Snakemake pipeline. This pipeline serves as an interface to the AlphaMissense community resource amino acid substitution dataset, which was made available upon the publication of AlphaMissense. By inputting a user-specified UniProt ID for any human protein in the AlphaMissense dataset, the pipeline extracts relevant substitution predictions, computes an average pathogenicity score per amino acid, and plots the results. This pipeline allows for easy customization to suit specific analysis goals for a target protein. Following the calculation of the average pathogenicity score per amino acid for ASNS using the Snakemake pipeline, further analysis was conducted to overlay these scores onto the human ASNS crystal structure. This overlay facilitated the visual identification of high pathogenicity scores in proximity to the two domain-specific substrate-bound active sites and the interface of the two domains, which is thought to encompass a critical intramolecular tunnel for ammonia transfer^7,28^. Residues associated with either active site or the tunnel residues were found to exhibit high predicted average pathogenicity scores. These results provided confidence in AlphaMissense’s calibration for ASNS and the high scores for the intramolecular tunnel residues further supported the recent report of the tunnel location within the protein structure^28^.

Given the concordance of AlphaMissense scores with known functional residues in ASNS, this study proceeded to assess AlphaMissense’s performance on established ASNSD-associated variants. Given that several recent reports have identified new ASNSD-associated variants, a comprehensive literature review was conducted to compile a list of known missense variants for evaluating AlphaMissense’s performance. The AlphaMissense classification results on these known variants indicated that it achieved a high classification accuracy for pathogenic variants. Notably, eight of the seemingly misclassified ASNSD variants classified as ambiguous or benign included five that were reported in patients with compound heterozygous conditions. These observations suggest that these variants may not necessarily be incorrectly classified by AlphaMissense, as their pathogenicity might be context-dependent, possibly requiring the presence of another pathogenic variant on the other allele, as would be the case in compound heterozygous patients. While this study did not undertake an analysis of combined scores for such ambiguous edge cases, future developments of AlphaMissense may enable researchers studying Mendelian diseases to do so. Additionally, we were unable to check the false positive rate of AlphaMissense due to the lack of a dataset of ASNS missense variants that can be confidently classified as benign. It is difficult to classify mutations as benign instead of unknown significance due to the homozygous and compound heterozygote genotypes in which ASNSD manifests, as well as the lack of an accurate diagnostic test to definitively rule out pathogenic effects.

The classification performance of AlphaMissense in identifying potential ASNSD-associated variants seems promising in this initial evaluation. These findings set the stage for further investigation into the significance of an AlphaMissense pathogenicity score for future ASNS variants. An examination would involve analyzing how a new variant fits into the broader distribution of pathogenicity scores and its specific location within the protein structure. It is important to clarify that this study’s primary focus is not to predict entirely new ASNS variants that might lead to ASNSD or, by itself, serve as a guide for prenatal genetic counseling. However, it was noted that a large number of novel pathogenic variants are predicted to be possible, covering a majority of all possible missense variants within the ASNS protein. In its current form, AlphaMissense should not be regarded as a replacement for the clinical assessment and experimental variant classification underlying a diagnosis of ASNSD. There is still much to be desired in the accuracy of computational tools for the prediction of variant effects in suspected cases of ASNSD and other diseases associated with missense mutations. However, the trends observed in this report could provide valuable insights, which, when combined with existing literature and other variant classification tools, may contribute to a timely estimation of the effect of new variants after they are detected by DNA sequencing. These collective efforts suggest that AlphaMissense holds potential as a useful tool for ASNS variant classification, potentially delivering meaningful diagnoses for patients and their families.

## Methods

### Snakemake pipeline

The Snakemake^29^ pipeline requires the user to specify a UniProt ID representing a protein of interest. Throughout its execution, the pipeline uses a predefined conda environment for each stage that is invoked automatically. All of the results generated by the pipeline are consolidated into a single directory, which includes final plots displaying the average AlphaMissense pathogenicity score for each amino acid in the specified protein and a heatmap of all scores for all variants. The Snakemake pipeline was created and verified for use on a Linux operating system and has not been validated on other operating systems.

### ASNSD literature review

We conducted a comprehensive review of ASNSD-related PubMed results with a specific focus on compiling known ASNS variants. The general variant classifications were maintained as per their original source references. Variants were renumbered as needed to account for discrepancies in reference sequence IDs. In ASNSD reports, it is common for variant positions to differ by one residue due to the unique characteristic of human ASNS having the first amino acid cleaved off before the final protein is formed^7^. A best effort has been made to preserve the original variant positions and accurately adjust positions that were different by one, based on the reference sequence data provided in the reports. In the literature review, all variants were listed, including those that either did not have a defined impact at the protein level (indicated in Table 1) or could not have their coordinate annotations resolved (only E360V). The variants with conflicts were not included in the processed list of missense ASNSD-associated variants used for benchmarking AlphaMissense.

### Analysis of ASNS variant predictions by AlphaMissense

The primary analysis of ASNS variant predictions by AlphaMissense was performed using the Snakemake pipeline. The critical residue positions overlayed onto the plot of average pathogenicity scores for all amino acid positions were obtained from the literature. The critical residues for the glutaminase N-terminal active site were defined as those within a hydrogen bonding network mediating substrate recognition and stabilization. The critical residues for the synthetase C-terminal active site were those defined by the overlay of bound AMP from the bacterial AS-B protein structure^8^. The critical tunnel residues were defined based on a recent report^28^. The exact coordinate system used for positioning critical residues was validated against the data for AlphaMissense. Custom scripts were used to sort through the ASNS substitution pathogenicity score results to obtain the variants of interest as related to ASNSD.

### Pymol

Using the graphical user interface of Pymol version 2.5.2^52^, the human ASNS structure was loaded using PDB structure 6GQ3^8^. Although the coloring process for the AlphaFold predicted structure is automated within the Snakamake pipeline, utilization of the known crystal structure was preferable to display bound substrates. All atoms were removed except for those in the first chain or the first chain’s bound substrates in the N and C terminal active sites. The structure was then colored by beta factors based on the average AlphaMissense pathogenicity score per residue as calculated using a custom script. The coloring process was performed by importing a custom Python script to the Pymol graphical user interface.

### Manuscript preparation

Figures were prepared using custom scripts, primarily utilizing the R packages ggplot2, cowplot, dplyr, grid, gridExtra, and png. Following an initial manually written draft of the manuscript, the AI-powered large language model ChatGPT 3.5 provided by OpenAI was used to refine the text for clarity.

## Acknowledgments

Funding for M.S.K. was from the National Institutes of Health, Institute of Child Health and Human Development (HD100576). S.J.S. is supported by a Starr Centennial Scholarship endowed to Cold Spring Harbor Laboratory from the Starr Foundation.

## Data Availability

The raw data underlying the analysis performed here was made publicly available as a community resource by the authors of the AlphaMissense paper. Results from the literature review on ASNSD are accessible in Table 1 and can be validated by accessing each paper by its provided PubMed ID (PMID) in the reference column of the table. The Snakemake pipeline and all data, figures, and the code to reproduce the analysis can be accessed on GitHub at https://github.com/StephenStaklinski/alphamissense_asns.

